# Nanoparticle Mediated NADPH Regeneration for Enhanced Ethanol Production by Engineered *Synechocystis* sp. PCC 6803

**DOI:** 10.1101/529420

**Authors:** Rajendran Velmurugan, Aran Incharoensakdi

## Abstract

The ethanol synthesis pathway engineered *Synechocystis* sp. PCC 6803 was used to investigate the influence of metal oxide mediated NADPH regeneration on ethanol synthesis. Among the metal oxides, Fe_2_O_3_ and MgO showed considerable improvement in growth, chlorophyll *a* content and ethanol synthesis. The *in-vitro* studies proved that the selected metal oxides have the potential to regenerate the NADPH under light illumination. The results clearly indicate that the light energy is the key factor for activation of metal oxides and to a less extent light itself has the possibility for direct regeneration of NADPH. Under optimized light intensity and NADP addition, the maximum MgO mediated ethanol production of 5100mg/L, about a 2-fold increase compared to the control, was obtained after 20 days cultivation at 5L level. This study indicates that the efficient NADPH regeneration aided by metal oxide is crucial to improve ethanol productivity in *Synechocystis* sp. PCC 6803.

**IMPORTANCE:** Cyanobacteria are efficient ethanol producing organisms from atmospheric CO_2_ upon engineering of pathway. In cyanobacterial ethanol synthesis pathway, NADPH plays an important role acetaldehyde to ethanol conversion. Here we elucidated the NADPH regeneration through extracellular addition of metal oxides. The metal oxide mediated NADPH regeneration study allows us to dissect the importance of metal oxides in enhancing ethanol production through NADPH regeneration while also providing insight into the regulatory functions of metal oxides in growth, photosynthetic apparatus and various carbon metabolisms.

---

To replace the fossil fuels, extensive studies have been focused on developing efficient techniquefor the production of ethanol from cyanobacteria (1,2). The incorporation of ethanol synthesis pathwayfrom various microorganisms into cyanobacteriahas already been demonstrated as a route of direct carbon fixation for ethanol production (3). However, the longercultivation time and lower yield at large scale makes the process ineffective (4). Apart from thepathway engineering strategies, there are certain factors such as cellular transport system, antioxidant enzymes, and cofactors influencing ethanol production and tolerance (5). Among them, NADPH plays an important role in photosynthesis, carbon metabolism, and more importantly NADPH plays an essential role in the conversion of acetaldehyde to ethanol (6). The engineering of various enzymes involved in NADPH regeneration to improve various functions of cyanobacteria has been demonstrated (6,7). On the other hand, the regeneration of NADPH is also reported as an effective process; however, it needs further study about activation of regenerating factor especially under photoautotrophic condition (8,9). Generally, the regeneration of cofactor (NADPH/NADH) has been performed in the presence of oxidoreductases (10) such as alcohol dehydrogenase (9), formate dehydrogenase (11), glucose dehydrogenase (12) and phosphite dehydrogenase (13); as well as redox molecules including 2-propanol, formate, glucose and glutamate (14); and light (8). In a recent study, the metal nanoparticles have also been highlighted as redox molecule for cofactor regeneration (14). The extracellular addition of metal nanoparticles at elevated concentration for regeneration of NADPH could be an economical process. However, the toxicity of metal nanoparticle and membrane mediated cellular intake of nanoparticle are the major concern for the successful process. The intake of metals by microalgae cells involves two mechanisms such as adsorption of metals on the cell surface containing functional groups (carboxyl, hydroxyl, phosphate, amino, sulfhydryl) and intake of metals through metal transport system (15). The metal transport system already exists naturally in case of *Synechocystis* and the identified metal transport related genes are *CtaA* and *PacS* (Cu) (16), *FutABC* transporter (Fe) (17), *MgtE* (Mg^2+^) (18), *mntABC* (Mn) (19), *ModD* and *ModC*(Mo) (18) and transporting P-type ATPase (Zn) (20). As a component of photosynthetic system, the metals are involved in various functions such as electron transport (Cu, Fe), PSII system (Mn), coordination of chlorophyll ring (Mg), water oxidation (Mn) and CO_2_ fixation (Zn). On the other hand, they are alsoinvolvedin various carbon metabolism along with the enzymes such as dinitrogenase reductase (Fe) (21), Mn-superoxide dismutase and Mn-catalase (Mn) (22), nitrate reductase (Mo) (23) and alcohol dehydrogenase /carbonic anhydrase (Zn) (24). In an attempt to analyse the influence of metal oxide nanoparticle on plant cells, researchers performed various studiesand the results showed that the metal oxide nanoparticle increases germination and growth by improving nitrate reductase activity in soybean (nano-SiO_2_ and nano-TiO_2_) (25); improves the growth of spinach by promoting photosynthesis and nitrogen metabolism (nano-TiO_2_) (26); increases the growth of spinach seeds by stimulating photosynthesis (TiO_2_ nanoparticles) (27) and improves the growth of *Cicer arietinum* and *Vignaradiata* (ZnO nanoparticles) (28).

The nanoparticles have been widely used to improve various cellular mechanisms for several decades. However, no study has been reported for the regeneration of NADPH under photoautotrophic condition. In the present study, the effect of metal oxides on growth and ethanol synthesis was analysed in *pdc-adh* overexpressing *Synechocystis*. The influence of selected metal oxides on NADPH regeneration was determined by *in-vitro* studies in which the most influential factors such as metal oxide concentration, light intensity and NADP concentration were studied in detail. Based on the *in-vitro* results, the cultivation media and environmental condition for *Synechocystis* were designed to limit thelevel at which metal, co-factor and light intensity exerts only beneficial effects on growth and biosynthesis of ethanol.

## MATERIALS AND METHODS

### Materials

The chemicals such as CuO (BDH Chemical Ltd, England), Fe_2_O_3_ (Sigma Aldrich), MgO (QReC), MnO (QReC), MoO_2_ (Sigma Aldrich), ZnO (Sigma Aldrich) and sulphuric acid (QReC) used in this study were analytical grade. The NADP, NADPH and all the standards used for high performance liquid chromatography (HPLC) analysis were the products of Sigma Aldrich.

### Microorganism and cultivation conditions

The *pdc-adh* pathway engineered *Synechocystis* sp. PCC 6803 (Pasteur Institute, France) was propagated on BG-11 agar medium (29). The expression vector pAPX was constructed by insertion of alcohol dehydrogenase (*adh* from *Synechocystis*) and pyruvate decarboxylase (*pdc* from *Saccharomyces cerevisiae*) genes into pEERM vector under the control of *psbA* promoter (unpublished). The engineered strain was cultivated in 250 mL Erlenmeyer flasks containing 100 mL BG-11 medium (pH 7.5) supplemented with 30 µg/mL chloramphenicol under continuous illumination of 100 μE/m^2^/s at 28±1°C with atmospheric CO_2_ supply upon shaking at 160 rpm.

### Screening of metal oxides for enhanced ethanol production

The screening of metal oxides such as CuO (1, 2 and 3 µM), Fe_2_O_3_ (10, 20 and 30 µM), MgO (100, 200 and 300 µM), MnO (10, 20 and 30 µM), MoO_3_ (1, 2 and 3 µM) and ZnO (1, 2 and 3 µM) were performed at various concentrations. The concentration ranges of metal oxides were obtained by comparing theconcentration of BG11 medium and inhibitory level (29,30,31). The effects of various metal oxides at various concentrations on biomass, chlorophyll *a* and ethanol concentration were determined. The engineered *Synechocystis* was cultured (initial OD≈0.06 at 730 nm) in 250 mL Erlenmeyer flasks containing 100 mL BG-11 medium (pH 7.5) with a continuous illumination of 100 μE/m^2^/s at 28±1 °C under atmospheric CO_2_ supply upon shaking at 160 rpm. The cells were harvested after 20 days of cultivation by centrifugation at 6000 × g for 10 min at room temperature. The supernatant was used for the analysis of extracellular products such as acetate, ethanol, pyruvate and succinate. For the analysis of intracellular products, the harvested cells were vigorously mixed with 500 μL of 70% methanol by vortex mixer. The mixture was incubated for 2 h at room temperature and then centrifuged at 6000 × g for 10 min at 4 °C. The supernatant was collected and dried in a vacuum evaporator at 40 °C. Pellet left after drying was dissolved and mixed thoroughly in 250 μL of water and 50 μL of chloroform followed by centrifugation at 6000 × g for 10 min (32). The uppermost water phase (200 μL) was collected, and filtered through a 0.45 μm Millipore filter before the analysis of acetate, ethanol, pyruvate and succinate by HPLC.

### In-vitro studies on NADPH regeneration

The effects of NADP, metal oxides, and light intensity on NADPH regeneration were studied by adding NADP (100 µM), alcohol dehydrogenase (200 U) and acetaldehyde (10 mM) along with Fe_2_O_3_(20 µM) or MgO(200 µM) in a total volume of 5 mL. Under optimized NADP and metal oxides, the samples were kept with a continuous illumination of 0, 50, 100, 150 and 200 μE/m^2^/s at 28±1 °C under shaking at 160 rpm. After 3 h of incubation, the samples were withdrawn and analysed for NADPH regeneration and ethanol production.

### In-situ studies on NADPH regeneration

The effect of metal oxide (Fe_2_O_3_:20 µM and MgO: 200 µM) mediated NADPH regeneration on biomass, chlorophyll *a* and ethanol production was analysed by adding the NADP (100 µM). The metal oxides such as Fe_2_O_3_ and MgO were used as NADPH activating materials. The effects of light/dark cycles were analysed by growing the cells in Erlenmeyer flask (250 mL) containing 100 mL BG-11 medium under dark and light (100 μE/m^2^/s) cycles for 20 days with 12 h alternate treatment. The effect of continuous light illuminationon ethanol production was studied by growing the cells in Erlenmeyer flask containing 100 mL BG-11 medium under light (100 μE/m^2^/s) for 20 days.

The scaleup experiment was performed in a 5 L photo-bioreactor equipped with controlled atmospheric air supply and the valves for inoculation and temperature monitor. The cells were cultivated in BG-11 medium with the atmospheric air supplementation at the flow rate of 200 mL/min under light (100 μE/m^2^/s) up to 25 days.

### Analytical methods

Intracellular pigments of *Synechocystis* cell suspension were extracted by dimethylformamide. Chlorophyll *a*was determined according to the method of Moran (33). The polyhydroxybutyrate (PHB) content was determined as described by Monshupanee and Incharoensakdi (34) using HPLC (Shimadzu, Japan) equipped with InertSustain 3-µm C18 column (GL Sciences, Japan) and UV/Vis detector. The estimation of lipid content was performed using the method described by Monshupanee and Incharoensakdi (34). The estimation of glycogen was performed by acid hydrolysis followed by sugar analysis by HPLC and the theoretical factor 1.111 was used for the glycogen to glucose conversion. The sugar and ethanol contents were quantified using HPLC system equipped with refractive index detector (RID 10A, Shimadzu, Japan). Metabolic intermediates such as acetate, pyruvate and succinate were quantified using HPLC equipped with UV/Vis detector (SPD-20A, Shimadzu, Japan). The components were separated in Phenomenex, Rezex ROA-Organic acid column (150 × 7.8 mm) with 5 mM H_2_SO_4_ as a mobile phase at a flow rate of 0.6 mL/min. The regeneration of NADPH was determined by the method described in Sigma Aldrich protocol (MAK038). Briefly, the samples were centrifuged at 10000 × g for 10 min and filtered to remove the protein. The filtered samples were used for total NADP (NADP and NADPH) and heated samples (60°C for 30 min) were used for NADPH analysis. Total NADP and NADPH samples were individually quantified at 450 nm on a plate reader.

### Statistical analysis

All experiments were performed in triplicate and the average values are reported. The average and standard deviation values were calculated using the respective functions (AVERAGE, STDEV) available in Microsoft Excel and the maximum difference among the three values was less than 5% of the mean.

## RESULTS AND DISCUSSION

### Screening of metal oxides for ethanol production

The effects of various metal oxides such as CuO, Fe_2_O_3_, MgO, MnO, MoO_2_ and ZnO on biomass, chlorophyll *a* and ethanol concentration were determined by cultivating the *pdc-adh* engineered *Synechocystis* in BG11 medium for 20 days. Among the metal oxides analysed, Fe_2_O_3_, MgO, MoO_3_ and ZnO showed positive effect on growth, while CuO and MnO showed inhibitory effect on growth (Fig. 1a). The oxidation states of Cu(I) and Cu(II) make the Cu an ideal factor for oxidoreduction reaction. However, they generate reactive oxygen species through Fenton and Haber-Weis reaction, thereby affecting the lipids, proteins and DNA (35). This nature of Cu affects the *Synechocystis*, thereby reducing the biomass concentration (Fig. 1a). Another metal, Mn is also acting as a cofactor for superoxide dismutase and catalase, butit reduces the growth (Fig. 1a) by competitively inhibiting the intake of other cofactors at higher concentration (36). As a cofactor, Mo plays an important role in nitrogen assimilation with the combination of Fe-Mo, where the higher Mo concentration interferes with the Fe balance inside the cell and further reduces the growth of cyanobacteria (37). The toxicity of nanoparticle and its severity are different among various organisms. Mahajan et al. (28) analysed the effect of nano-ZnO on the growth of *Vigna radiate* and *Cicer arietinum* and observed maximum tolerable level of 20 ppm and 1 ppm, respectively. In another study, the effect of CuO nanoparticles on *Microcystis aeruginosa* was analysed and the smaller nanoparticles could enter the cells and form excess ROS, damage DNA, affect the membrane integrity and finally affect the growth (30). As mentioned by Auffan et al. (38), dissolution might be another factor in nanoparticle toxicity by impacting the release of undesired ions, which can further stimulate the formation of radicals in the presence of light. The results clearly indicate that the nanoparticles are coherent to form superoxides; however, the metal nanoparticles are not always negatively affecting the cell growth. The highest biomass concentration was observed when treated with F_*2*_O_3_, MgO and ZnO, which denotes the positive effect of metal oxides on the growth (Fig. 1a). The positive effect of Fe_2_O_3_, MgO and ZnO is well documented for *Synechocystis* where they play an important role in photosynthesis and nitrogen fixation (39).

**FIG 1.**
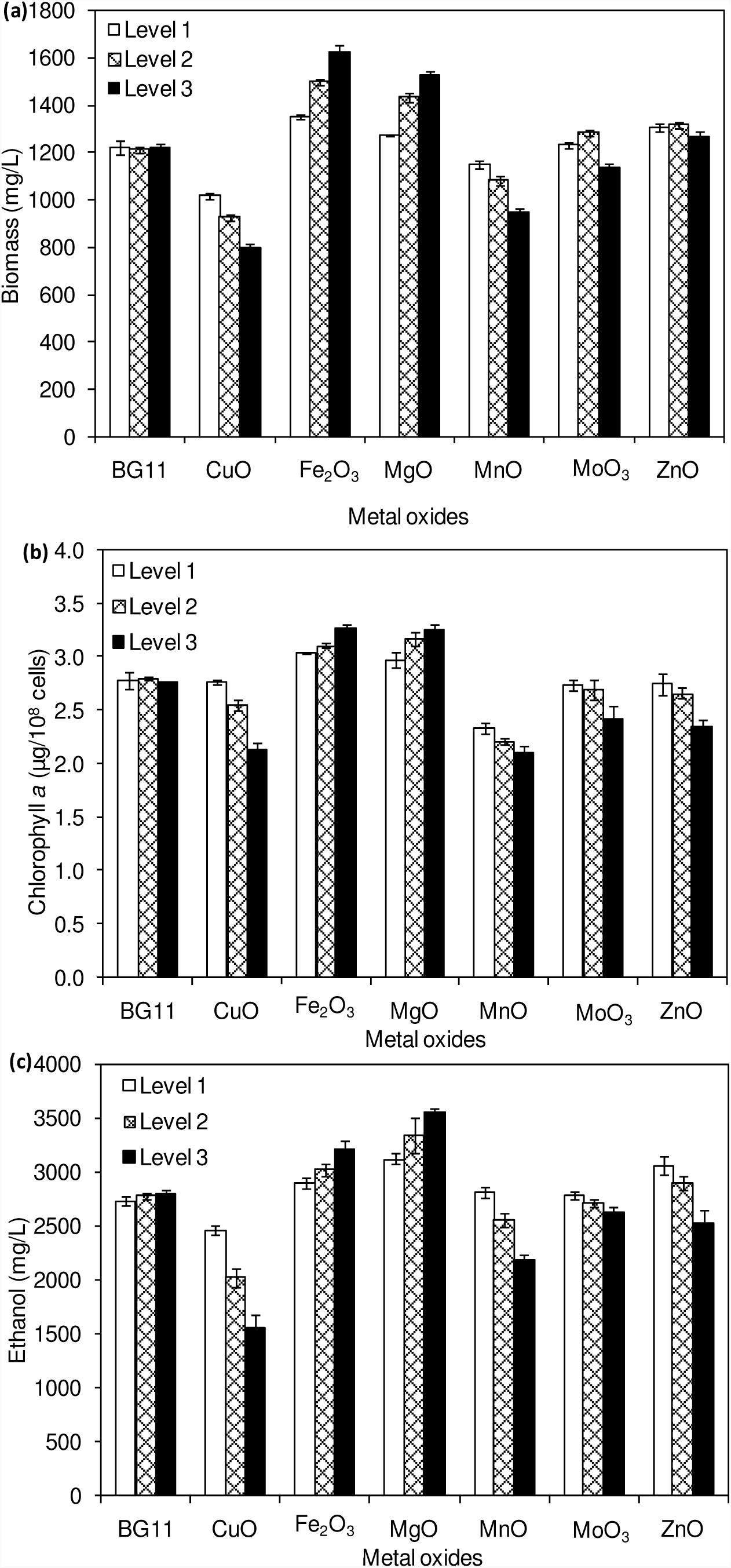
Effect of metal oxides at various concentrations on biomass (a), chlorophyll *a* (b) and ethanol (c) concentration (conditions: cultivation time: 20 days; CuO (1, 2 and 3 µM), Fe_2_O_3_ (10, 20 and 30 µM), MgO (100, 200 and 300 µM), MnO (10, 20 and 30 µM), MoO_3_ (1, 2 and 3 µM) and ZnO (1, 2 and 3 µM).

The chlorophyll*a* content was increased upon an increase in Fe_2_O_3_ and MgO concentration, whereas MoO_3_ and ZnO reduced chlorophyll *a* slightly upon an increase in concentration (Fig. 1b). In contrast, CuO and MnO decreased the chlorophyll *a* content drastically upon an increase in concentration. The increase in chlorophyll *a* content by Fe_2_O_3_ and MgO is due to the functional properties of these metals in chlorophyll *a* (39). It is reported that the iron limitation in algae/cyanobacteria results in reduced levels of cytochrome *f* (related to chlorophyll) and phycobilin pigment (40).On the other hand, the CuO, MnO, MoO_3_ and ZnO reduced the chlorophyll *a* content with a subsequent decreased biomass concentration. The decrease of ethanol concentration by CuO and MnO is due to their toxicity to *Synechocysits*; however, the cofactor activity of Mn in pyruvate carboxylase catalysed reaction might be another factor, which causes direct utilization of pyruvate for oxaloacetate formation (41). Although the MgO showed comparatively lower growth with respect to Fe_2_O_3_, the ethanol concentration was high with MgO which indicates its favorable contribution towards ethanol production (Fig. 1c). The storage carbon such as glycogen, PHB and lipid contents were analysed to understand the flux of carbon sources. As can be seen in Fig. 2, the Fe_2_O_3_ and MgO increased the glycogen, PHB and lipid contents. The MoO_3_ increased the contents of glycogen and PHB, while the CuO increased the lipid accumulation.

**FIG 2.**
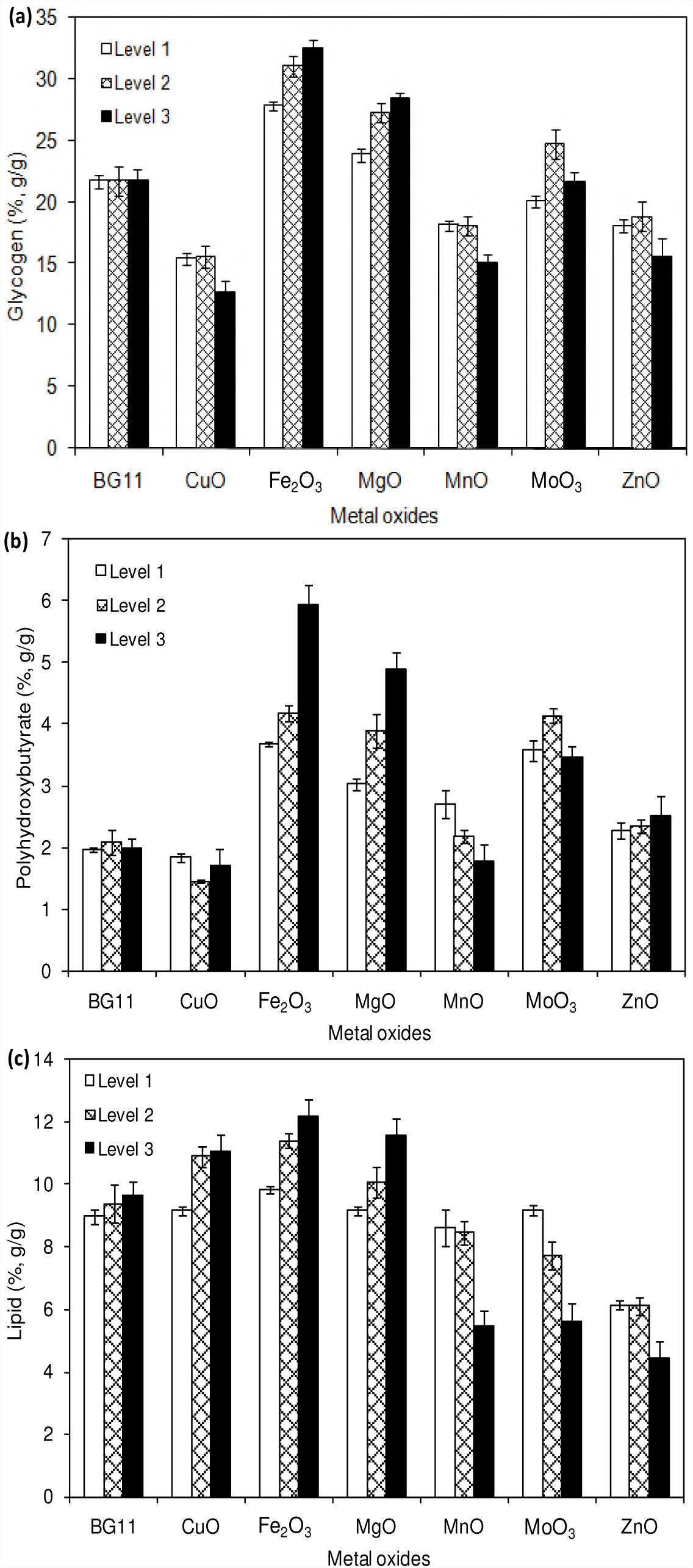
Effect of metal oxides at various concentrations on glycogen (a), polyhydroxybutyrate (b) and lipid concentration (conditions: cultivation time: 20 days; CuO (1, 2 and 3 µM), Fe_2_O_3_ (10, 20 and 30 µM), MgO (100, 200 and 300 µM), MnO (10, 20 and 30 µM), MoO_3_ (1, 2 and 3 µM) and ZnO (1, 2 and 3 µM).

### In-vitro studies on NADPH regeneration in the presence of light and metal oxides

The integrated chemical/photo-enzymatic regeneration of NADPH is the process of oxidation of NADP in oxidoreductase mediated enzymatic reaction in the presence of oxidizing molecule and light energy. To analyse the effect of metaloxides andlight intensity on NADPH regeneration, the reaction was integrated with alcohol dehydrogenase catalysed conversion of acetaldehyde to ethanol. As can be seen in Fig. 3a, each metal oxideshowed its uniqueactivity on NADPH regeneration. Compared to the control (BG11 media or water), all the metal oxides showed higher regeneration potential. The maximum regeneration was observed with MgO followed by Fe_2_O_3_, CuO, MnO, ZnO and the least NADPH regeneration was observed with MoO_3_. The variation in regeneration might be ascribed to the interface, structural and electronic properties of individual metal oxides (Picone et al., 2016) (42). Similar to the present study, Lee et al. (43) used SiO_2_, glutamate dehydrogenase and visible light for the regeneration of NADH, in which the authors observed 100% conversion of α-ketoglutarate to glutamate after 5 h of treatment. To reduce the dependency on enzymes, researchers focused on regeneration of NADPH in the absence of enzyme by varying the metal/metal oxidessuch as Rh, SiO_2_ and TiO_2_and observed significant improvement in regeneration (44,45). To analyse the photochemical regeneration of NADH, Jiang et al. (44) used carbon-containing TiO_2_ and reported 63.98% conversion under visible light. However, those metal oxides reported for cofactor regeneration may also cause some hazardous effect on cyanobacteria (46). The compatibility of metal oxide is a major concern to perform in-situ process development. In this study, the type and concentration of metal oxides were chosen based on the metals present in the cyanobacterial medium (29) and the influences of metal oxide concentration were studied. As can be seen in Fig. 3a, the metal oxides used were compatible with cyanobacteria, in which MgO showed highest NADPH regeneration activity followed by that with Fe_2_O_3_. Thus, the effect of light intensity on NADPH regeneration was analysed for MgO and Fe_2_O_3_ by varying its ranges from 50to 200 µE/m^2^/s. The result showed that an increase in light intensity constantly increased the NADPH regeneration (Fig. 3b). However, when comparing two metal oxides, the MgO always showed higher regeneration. The control experiment in the absence of light also regenerated the NADPH upto a certain level, which might be due to the reactivity of metal oxides to regenerate the NADPH. Factors such as metal oxide and light illumination may also cause the harmful effect on cyanobacteria, thus, it is important to analyse its compatibility in *in-situ* experiments.

**FIG 3.**
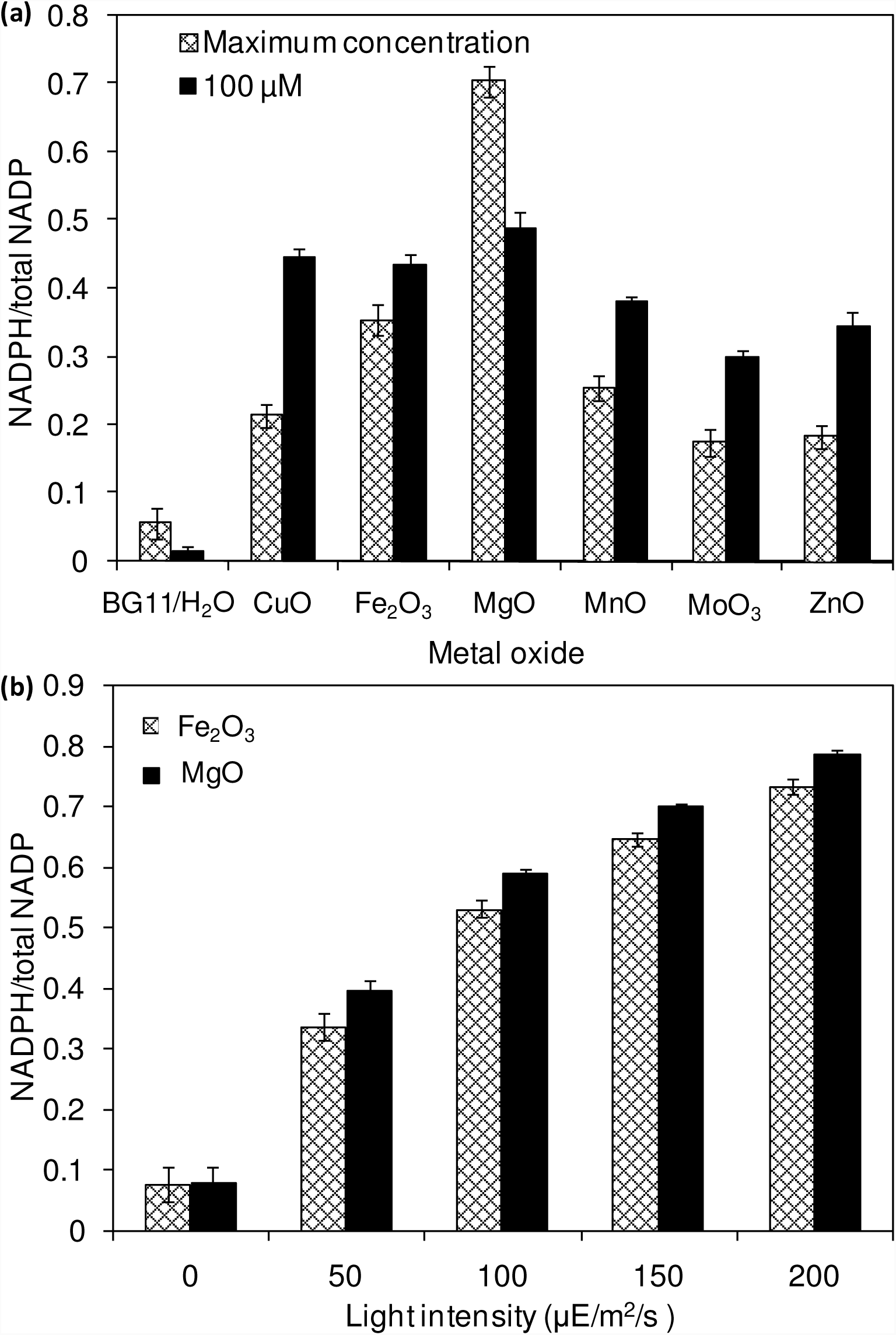
*In-vitro* analysis on influence of metal oxides on NADPH regeneration (conditions: cultivation time: 20 days; CuO (2 µM), Fe_2_O_3_ (20 µM), MgO (200 µM), MnO (20 µM), MoO_3_ (2 µM) and ZnO (2 µM) (a), and light intensity on Fe_2_O_3_ (20 µM) and MgO (200 µM) mediated NADPH regeneration (b).

### Effect of NADP concentration on biomass, chlorophyll *a* and ethanol production

As presented in Fig. 4, the effect of NADP concentration on biomass, chlorophyll *a* and ethanol concentration were analysed by varying the concentration from 50 to 250 µM. The increase in NADP increased the biomass and chlorophyll *a* in all the experiments such as control (without metal oxide), with Fe_2_O_3_ and MgO. Similarly, an increase in NADP concentration increased both intracellular and extracellular acetate, succinate, pyruvate and ethanol concentrations (Table 1). The control experiments (without metal oxides) showed higher extracellular acetate, succinate and pyruvate, and lower ethanol content, while the metal oxide addition increased the ethanol content but reduced the other metabolites. The results clearly indicate the redirection of metabolites of very closely related pathway into ethanol synthesis mediated by metal oxides, and the highest ethanol was observed with150 µM NADP (Table 1). Thus, metal oxides facilitate the reduction of added NADP into NADPH, which is further used for carbon fixation and biosynthesis topromote the growth of *Synechocystis* (47). Recently, there was an attempt to engineer glucose-6-phosphate dehydrogenase gene (pentose phosphate pathway) in *Synechocystis* sp. PCC 6803 to enhance the reduction of NADP into NADPH. As a result of improved NADPH level intracellularly, the biomass and ethanol concentrations were increased (6). In the present study, the increased NADPH regeneration mediated by metal oxides has been achieved without engineering other natural pathways of *Synechocystis* and the production of ethanol is also higher than that reported by Choi and Park (6).

**Table 1.** Effect of NADP concentration on intracellular and extracellular products in three different conditions.

| NADP<br>(μM) | Intracellular (mg/g) | | | | Extracellular (mg/L) | | | |
| --- | --- | --- | --- | --- | --- | --- | --- | --- |
|  | Acetate | Succinate | Pyruvate | Ethanol | Acetate | Succinate | Pyruvate | Ethanol |
| Control (no Fe<sub>2</sub>O<sub>3</sub> / MgO) |  |  |  |  |  |  |  |  |
| 0 | 3.8±0.4 | 2.9±1.1 | 4.2±1.1 | 11.3±1.5 | 121.2±8.1 | 116.3±7.6 | 68.2±9.8 | 2822.0±245.7 |
| 50 | 4.6±0.8 | 4.1±0.7 | 6.7±0.8 | 13.9±0.9 | 151.7±10.7 | 143.5±10.2 | 94.1±11.7 | 3076.2±184.1 |
| 100 | 9.5±0.6 | 8.6±0.6 | 10.9±0.3 | 16.4±0.7 | 168.1±11.6 | 169.5±8.1 | 115.5±11.2 | 3268.6±214.0 |
| 150 | 11.7±1.0 | 11.0±0.7 | 13.3±1.0 | 17.6±1.2 | 180.6±11.7 | 186.4±6.5 | 121.3±9.7 | 3391.0±278.3 |
| 200 | 12.3±0.8 | 11.4±0.9 | 13.5±1.1 | 18.4±1.1 | 183.7±12.5 | 192.1±5.9 | 130.7±10.4 | 3425.4±302.8 |
| 250 | 12.2±0.9 | 11.5±1.0 | 13.4±0.8 | 18.7±1.6 | 186.6±10.6 | 199.1±7.8 | 135.7±12.2 | 3511.2±310.4 |
| Fe<sub>2</sub>O<sub>3</sub> (20 μM) | | | | | | | | |
| 0 | 3.9±1.3 | 4.2±0.8 | 5.8±0.8 | 8.1±0.5 | 112.3±7.5 | 111.8±5.4 | 60.6±7.6 | 3013.3±235.6 |
| 50 | 5.4±0.4 | 5.0±0.2 | 7.4±0.5 | 10.9±0.7 | 136.4±10.4 | 132.1±7.0 | 84.4±9.2 | 3521.4±243.7 |
| 100 | 7.9±0.8 | 7.2±0.4 | 10.2±0.2 | 13.6±0.8 | 148.5±8.6 | 155.1±5.9 | 107.4±10.0 | 4138.8±243.0 |
| 150 | 10.6±0.9 | 9.1±0.9 | 11.3±0.4 | 15.7±0.4 | 162.0±11.3 | 163.0±5.8 | 112.1±12.4 | 4721.3±318.4 |
| 200 | 10.3±1.1 | 9.5±0.7 | 12.0±0.8 | 16.9±0.6 | 165.2±12.8 | 170.0±6.8 | 119.1±10.7 | 4746.8±374.9 |
| 250 | 10.5±0.9 | 9.8±0.6 | 12.1±0.7 | 16.8±0.9 | 160.2±11.6 | 172.8±4.6 | 126.6±11.8 | 4731.7±365.2 |
| MgO (200 μM) | | | | | | | | |
| 0 | 3.8±0.5 | 3.6±0.7 | 5.1±0.6 | 7.5±0.6 | 84.5±8.2 | 101.9±6.0 | 43.2±4.7 | 3752.3±270.5 |
| 50 | 5.5±0.6 | 5.4±0.3 | 6.9±0.8 | 10.2±0.7 | 98.5±8.7 | 119.1±5.1 | 63.5±6.8 | 4065.6±212.8 |
| 100 | 7.4±1.7 | 6.9±1.1 | 8.8±1.1 | 12.5±0.9 | 107.4±8.1 | 135.2±5.6 | 69.7±7.3 | 4721.3±198.3 |
| 150 | 11.8±1.1 | 11.0±0.7 | 13.3±1.0 | 17.6±1.5 | 120.2±16.4 | 145.0±5.9 | 78.3±8.2 | 4973.1±282.9 |
| 200 | 9.1±1.2 | 8.1±0.7 | 10.7±1.2 | 16.1±0.5 | 123.2±7.8 | 156.9±6.5 | 84.8±7.6 | 5028.1±216.0 |
| 250 | 9.4±1.1 | 8.4±0.9 | 11.0±0.4 | 16.4±0.6 | 123.6±5.5 | 154.4±10.4 | 88.2±7.4 | 5014.7±261.3 |

5 The values are triplicates of three different experiments and the ± values are standard deviation.

**FIG 4.**
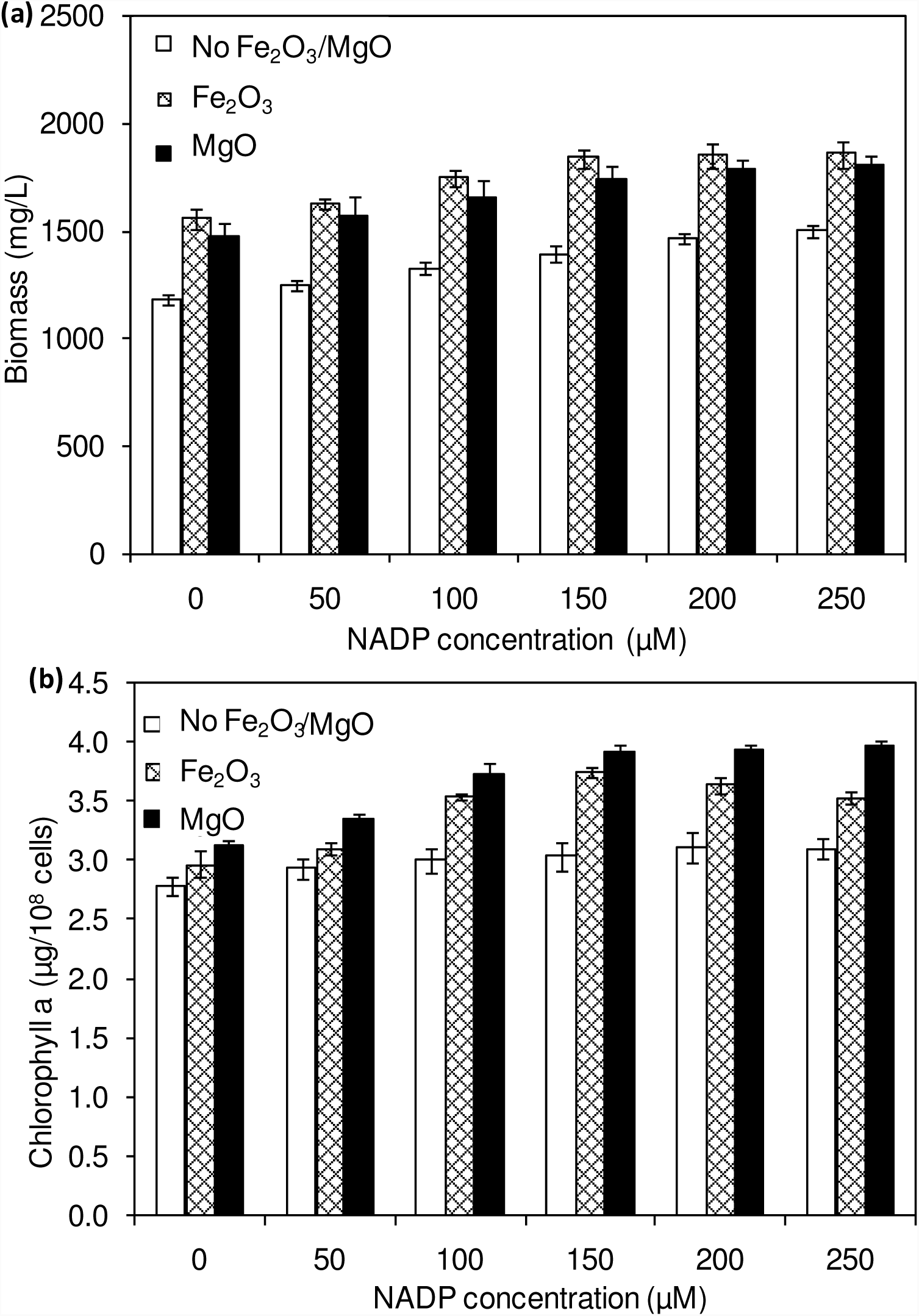
Effect of NADP concentration on biomass (a) and chlorophyll *a* (b) concentrations (conditions: cultivation time: 20 days; Fe_2_O_3_: 20 µM and MgO:200 µM).

### Effect of light intensity on biomass, chlorophyll a and ethanol production

The light acts as a strong metabolic valve for the photosynthetic organisms. In this context, we examined the influence of continuous light at various intensities on biomass, chlorophyll *a* and ethanol production (Fig. 5). The continuous light highly influencesbiomass, chlorophyll *a* and ethanol production, which signifies that the light illumination is an indispensable factor to sustain the growth rate and to regenerate the NADPH (9). It should be noted that even in the presence of externally added metal oxides and NADP, the continuous light does not affect the growth upto normal light illumination (100 µE/m^2^/s), which indicates the compatibility of the integrated treatment in cyanobacterial system (Fig. 5a). When comparing the metal oxides, the Fe_2_O_3_ showed higher biomass and chlorophyll *a* content, whereas the MgO produced higher ethanol production (Fig. 5 and Table 2). When comparing the light intensities, the light intensity higher than 100 µE/m^2^/s reduced the biomass, chlorophyll *a*content and ethanol production. Normally, the light intensity of 150 µE/m^2^/s does not cause harmful effect; however, the added metal oxides may reduce the growth by generating the excess free radicals. Both theintracellular andextracellular acetate, succinate, pyruvate and ethanol contentswere significantly increased upon an increase in light intensity upto 100 µE/m^2^/s, above which resulted in decreased contents (Table 2). Together, high biomass and chlorophyll *a* contents with continuous light at100 µE/m^2^/s intensity produced highest ethanol content of 4721 (Fe_2_O_3_) and 4973(MgO) mg/L.

**Table 2.** Effect of light intensity on intracellular and extracellular products in three different conditions.

| Light<br>($\mu\text{E}/\text{m}^2/\text{s}$) | Intracellular (mg/g) | | | | Extracellular (mg/L) | | | |
| --- | --- | --- | --- | --- | --- | --- | --- | --- |
|  | Acetate | Succinate | Pyruvate | Ethanol | Acetate | Succinate | Pyruvate | Ethanol |
| Fe<sub>2</sub>O<sub>3</sub>(20 $\mu\text{M}$) | | | | | | | | |
| 0 | 3.4±0.8 | 2.8±0.6 | 4.5±1.2 | 6.1±1.1 | 29.4±9.6 | 29.0±5.4 | 28.9±3.9 | 164.6±63.0 |
| 50 | 5.7±0.3 | 4.9±0.2 | 7.2±0.7 | 10.2±1.3 | 114.3±2.8 | 113.7±7.7 | 70.7±8.3 | 3102.7±150.7 |
| 100 | 10.6±0.9 | 9.1±0.9 | 11.3±0.4 | 15.7±0.4 | 162.0±11.3 | 163.0±5.8 | 112.1±12.4 | 4721.3±318.4 |
| 150 | 6.7±0.2 | 5.8±0.6 | 7.6±0.5 | 11.8±0.7 | 132.3±10.5 | 134.6±14.2 | 86.0±6.1 | 4534.1±293.0 |
| 200 | 6.3±0.5 | 5.8±1.1 | 7.1±1.6 | 11.7±1.4 | 122.1±12.7 | 125.2±22.3 | 79.6±5.5 | 3762.8±272.3 |
| MgO (200 $\mu\text{M}$) | | | | | | | | |
| 0 | 2.9±0.9 | 2.0±0.8 | 3.5±1.3 | 10.0±1.6 | 29.6±6.2 | 32.9±6.7 | 19.5±8.6 | 195.0±97.1 |
| 50 | 4.1±0.5 | 3.5±0.5 | 6.2±0.7 | 13.8±1.1 | 83.4±9.8 | 102.8±5.6 | 50.6±9.0 | 3246.7±271.5 |
| 100 | 11.8±1.1 | 11.0±0.7 | 13.3±1.0 | 17.6±1.5 | 120.2±16.4 | 145.0±5.9 | 78.3±8.2 | 4973.1±282.9 |
| 150 | 5.0±1.0 | 4.1±0.6 | 6.3±0.6 | 14.9±1.3 | 106.6±12.0 | 128.1±5.1 | 63.7±7.7 | 4908.5±224.4 |
| 200 | 4.7±0.7 | 3.7±0.4 | 5.6±0.8 | 12.0±0.9 | 99.3±10.7 | 120.5±6.0 | 57.5±7.5 | 4132.6±225.4 |

The values are triplicates of three different experiments and the ± values are standard deviation.

**FIG 5.**
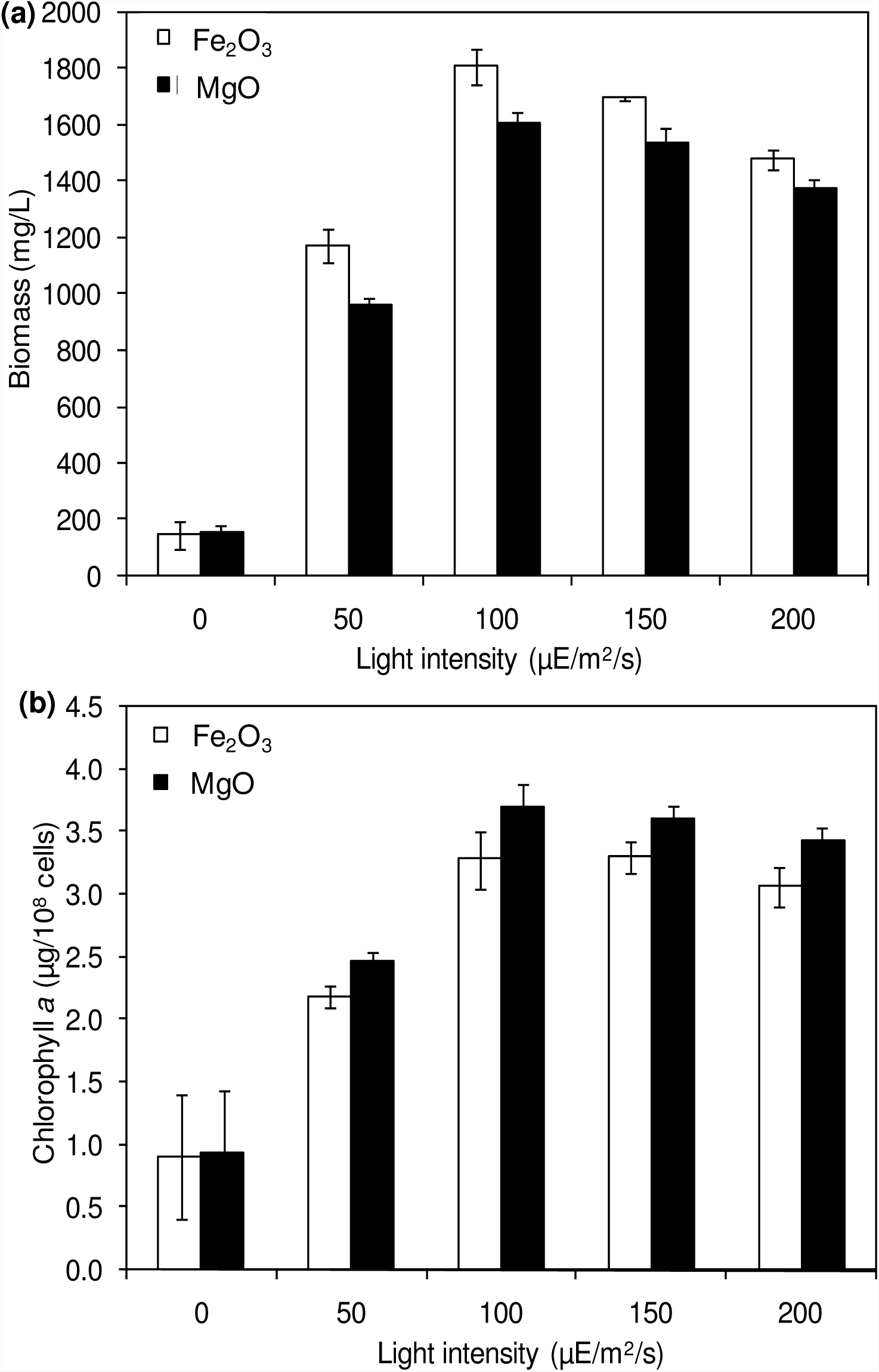
Effect of light intensity on biomass (a) and chlorophyll *a*(b) concentrations (conditions: cultivation time: 20 days; NADP: 150 µM; Fe_2_O_3_: 20 µM and MgO: 200 µM).

### NADPH regeneration facilitates ethanol production

The ethanol production was performed at 5 L level to analyse the feasibility of NADPH regeneration at large scale. As can be seen in Fig. 6a, the ethanol and biomass production was increased with an increase in incubation time and reached the maximum at 20 days. When comparing Fe2O_3_ and MgO with respect to ethanol production, the MgO showed higher ethanol productionsimilar to that observed in flask experiments. The ethanol levels with Fe_2_O_3_ and MgO treatments, at 4851 and 5100 mg/L respectively, were also very close to those of the flask experiments. The results clearly indicate the proper distribution of metal oxides (Fe_2_O_3_ and MgO), NADP and light are important for an optimalregeneration of NADPH in *Synechocystis* at large scale. The successful use of light illumination for co-factor regeneration has been previously reported which utilizes microorganisms as a source of the enzyme oxidoreductases (8,9).The alcohol dehydrogenase engineered *Rhodobacter sphaeroides* could regenerate the co-factor efficiently in the presence of light during the production of chlorophenyl ethanol (9). However, the present study showed for the first time the production of ethanol integrated with nanoparticle mediated NADPH regeneration.

**FIG 6.**
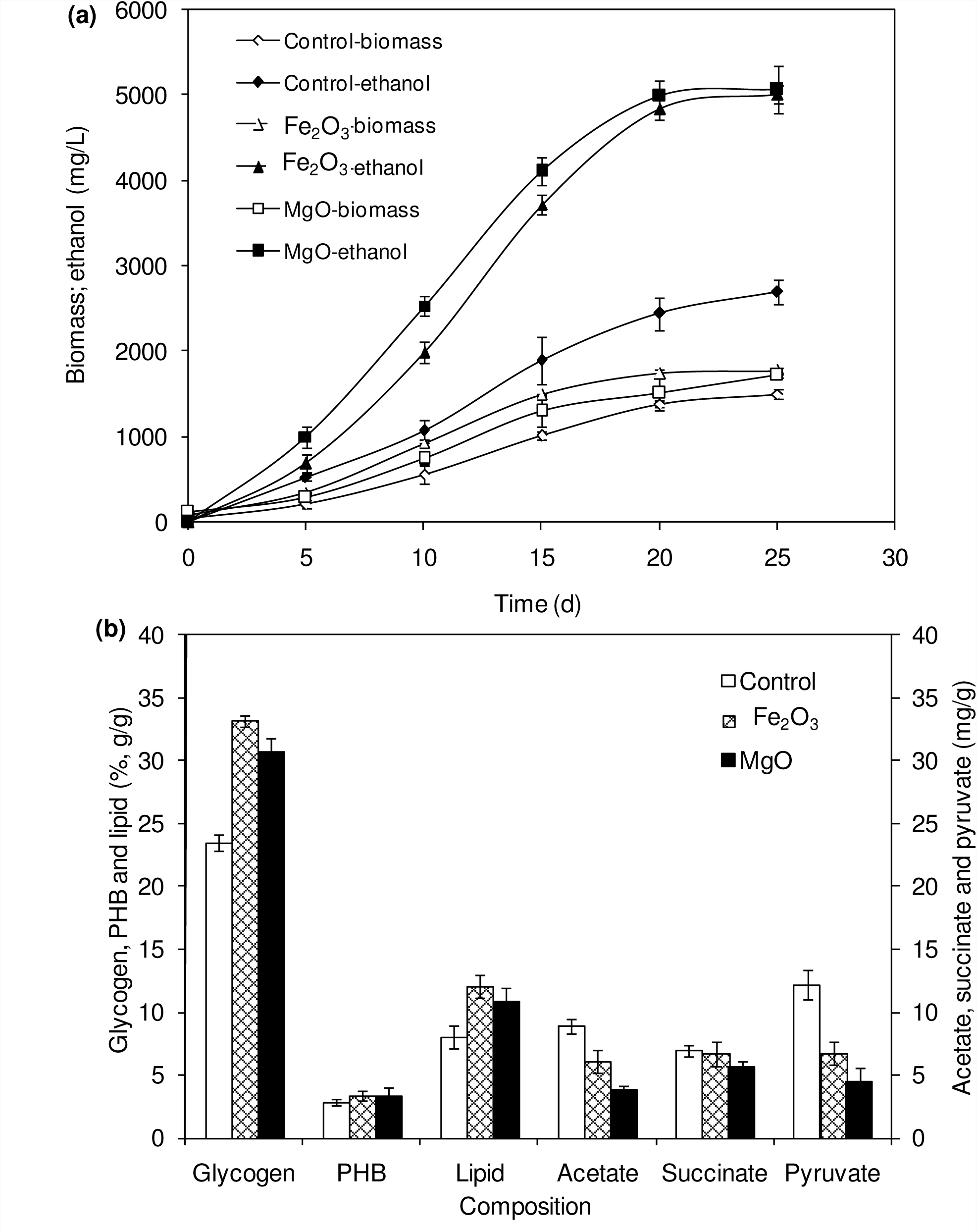
Effect of incubation time on ethanol and biomass concentration (a) and composition of intracellular products of *Synechocystis* (conditions: Fe_2_O_3_: 20 µM; MgO: 200 µM and NADP: 150 µM; light: 100 µE/m^2^/s).

### Flux in carbon metabolism upon NADPH regeneration

The intracellular glycogen, PHB and lipid contents in *Synechocystis* were analysed to monitor the changes in metabolism of engineered strain under NADPH regenerationat 5 L level (Fig. 6b). After 20 days of growth, the engineered strain produced 23.8% (g/g DCW) glycogen in BG11 medium containing 150 µM NADP, whereas it was increased upon the addition of Fe_2_O_3_ and MgO. The maximum glycogen content of 33.2% was observed with Fe_2_O_3_, whereas MgO produced 30.7%. The PHB content was slightly increased upon Fe_2_O_3_ and MgO addition; however, it is not upto considerable level. As observed with glycogen content, the lipid content also increased with Fe_2_O_3_ and MgO, whereas the maximum lipid content was observed with Fe_2_O_3_. The control values were always higher than that of metal oxide treatment without NADP, which indicates the NADP also influences glycogen, PHB and lipid contents (Fig. 2 and 6b). The 1.5-fold increase in lipid content upon NADPH regeneration was already reported by Osada et al. (48). The results further confirm that the presence of metal oxides regenerated the NADPH and improved the lipid accumulation.

The analysis of intracellular metabolites such as acetate, succinate and pyruvate showed the significant improvement in concentration upon NADPH regeneration (Fig. 6b). The highest intracellular acetate, succinate and pyruvate concentration of about 8.9, 7.0 and 12.2 mg/g, respectively was observed in the control experiment with NADP (no metal oxides). The addition of metal oxides reduced theintracellular acetate, succinate and pyruvate concentration considerably. The reduction in pyruvate content is likely due to the utilization of pyruvate for ethanol synthesis through acetaldehyde formation. Acetate is another metabolic intermediate for ethanol synthesis which is also acting as a substrate for ethanol synthesis (3). Succinate, the TCA cycle intermediate, was also reduced which might be due to the changes in NADPH/NADP ratio, as suggested by You et al. (47).

The *in-vitro* and *in-situ* results clearly demonstrate that the metal nanoparticles have the potential to regenerate the NADPH and their use in cyanobacterial system can regenerate the NADPH even in the absence of external NADP addition. The carbon flux in engineered *Synechocystis* metabolism indicates that the NADPH regeneration potentially redirects the carbon from very closely related pathway into ethanol synthesis pathway.

## CONCLUSION

As the cyanobacterial ethanol production involves phototrophic growth condition, the use of light energy for NADPH regeneration is feasible. However, the light energy alone has no potential to obtain an effective regeneration process. In these aspects, the metal oxides were applied in both *in-vitro* as well as *in-situ* experiments. The addition of metal oxides Fe_2_O_3_ and MgO significantly improved the NADPH regeneration, which further improved the ethanol production upto 5100 mg/L at 5 L experiments. Thus, we demonstrated that introducing nanoparticle mediated NADPH regenerationisa promising strategy to increase the ethanol production in engineered *Synechocystis*.

## ACKNOWLEDGEMENTS

R.V. is thankful to the Graduate School and Faculty of Science, Chulalongkorn University (CU), for post-doctoral fellowship from Ratchadaphiseksomphot Endowment Fund. A.I. acknowledges the research grant from CU on the Frontier Research Energy Cluster (CU-59-048 –EN) and from the Thailand Research Fund (IRG5780008).

## List of Tables

**Table 1** Effect of light intensity on intracellular and extracellular products under three different conditions.

**Table 2** Effect of NADP concentration on intracellular and extracellular products under three different conditions.

